# PPTC7 acts as an essential co-factor of the SCF^FBXL4^ ubiquitin ligase complex to restrict BNIP3/BNIP3L-dependent mitophagy

**DOI:** 10.1101/2024.06.20.599818

**Authors:** Xiayun Xu, Yingji Chen, Yao Li, Shi-Min Zhao, Chenji Wang

## Abstract

Mitophagy is a selective process that targets damaged, dysfunctional, or superfluous mitochondria for degradation through autophagy. The SCF^FBXL4^ ubiquitin ligase complex suppresses basal mitophagy by targeting BNIP3 and BNIP3L, two key mitophagy cargo receptors, for ubiquitin-proteasomal degradation. *FBXL4* loss-of-function mutations lead to excessive BNIP3/3L-dependent mitophagy, thereby causing a devasting multi-system disorder called mitochondrial DNA depletion syndrome, type 13 (MTDPS13). PPTC7, a mitochondrial matrix phosphatase, is essential for proper mitochondrial function and biogenesis. Here, we show that a proportion of PPTC7 is located on the outer mitochondrial membrane, where it interacts with FBXL4 and BNIP3/3L. PPTC7 decreases BNIP3/3L protein stability in a protein phosphatase activity-independent manner. Using *in vitro* cell culture and *Pptc7* knockout mice models, we demonstrate that PPTC7 deficiency activates high levels of basal mitophagy in a BNIP3/3L-dependent manner. Mechanistically, PPTC7 facilitates SCF^FBXL4^-mediated ubiquitin-proteasomal degradation of BNIP3/3L. Overall, these findings establish PPTC7 as an essential co-factor of the SCF^FBXL4^ complex and a suppressor of BNIP3/3L-dependent mitophagy.

## INTRODUCTION

Mitochondria are essential organelles in eukaryotic cells, playing vital roles in crucial cellular processes such as bioenergetics, metabolism, and signal transduction. Mitophagy is a cellular process involving the targeted degradation and removal of damaged, dysfunctional, or superfluous mitochondria by autophagy, essential for maintaining cellular homeostasis and mitochondrial quality control. This selective removal of impaired mitochondria helps to prevent the accumulation of defective organelles, thereby reducing oxidative stress, maintaining energy production balance, and promoting cellular homeostasis^1–3^. Dysregulation of mitophagy has been linked to a wide range of conditions, including neurodegenerative disorders, metabolic disorders, as well as cancer and cardiovascular diseases. Impaired mitophagy can lead to the buildup of damaged mitochondria, resulting in increased oxidative damage, inflammation, energy deficits, and ultimately contributing to disease progression and pathology^2, 4^. Conversely, excessive mitophagy under certain pathological conditions can decrease mitochondrial content, burdening the remaining organelles and ultimately triggering mitophagic cell death^5, 6^.

Most studies on mitophagy have focused on a canonical pathway involving Parkinson’s disease-related proteins PINK1 and Parkin, particularly their roles in depolarization-induced mitophagy *in vitro*^7, 8^. However, recent research using Mito-QC reporter in mice and Drosophila models revealed that basal mitophagy activity is largely independent of the PINK1/Parkin pathway^9^. Alternatively, mitochondrial outer membrane receptors, including but not limited to BNIP3, BNIP3L/NIX, FUNDC1, and BCL2L13, also play crucial roles in mitophagy^4, 10^. These receptors serve as bridges between the mitochondria marked for degradation and the autophagic machinery, ensuring efficient removal of these organelles. By interacting with ATG8 family proteins (LC3/GABARAPs) on the autophagosomal membrane, these receptors help trigger the formation of autophagosomes around the targeted mitochondria, leading to their subsequent degradation^11^. Among them, BNIP3 and BNIP3L are transcriptionally activated by the transcription factor hypoxia-inducible factor (HIF1α) under hypoxia, thereby promoting cellular adaptation to low-oxygen conditions and ensuring efficient metabolic remodeling for cell survival^12^.

We and others have recently demonstrated that the ubiquitin-proteasomal degradation of BNIP3 and BNIP3L are tightly regulated by a SCF^FBXL4^ E3 ubiquitin ligase complex^13–16^. FBXL4, a member of the F-box protein family, serves as a receptor for substrates to facilitate their recognition by the Skp1-Cul1-F-box (SCF) E3 ubiquitin ligase complex. Importantly, biallelic mutation in the *FBXL4* gene lead to Encephalomyopathic mitochondrial DNA (mtDNA) depletion syndrome 13 (MTDPS13), a severe infantile-onset genetic disorder characterized by excessive mitophagy in patient tissues and organs^17–19^. MTDPS13-associated FBXL4 mutations disrupt the assembly of an active SCF^FBXL4^ complex, resulting in the robust accumulation of BNIP3/3L proteins, triggering high levels of mitophagy even under basal conditions, underscoring the harmful effects of excessive mitophagy on cellular homeostasis^13–16^.

Although the pathophysiological roles of SCF^FBXL4^ complex in mitophagy have been clearly established, little is known about how this complex is regulated. Here, we demonstrate that PPTC7, a mitochondrial matrix phosphatase^20^, interacts with FBXL4 and BNIP3/3L on the mitochondrial outer membrane. PPTC7 deficiency leads to excessive mitophagy in a BNIP3/3L-dependennt manner but independent of its protein phosphatase activity. We establish PPTC7 as an essential co-factor for the SCF^FBXL4^ E3 complex to facilitate ubiquitination and degradation of BNIP3/3L.

## RESULTS

### Identification of BNIP3/3L and FBXL4 as PPTC7-interacting proteins

To elucidate the unidentified functional partners of FBXL4, we analyzed the genetic co-dependency between FBXL4 and other proteins using Broad’s 21Q2 DepMap dataset^21^. This dataset, derived from large-scale loss-of-function sgRNA screens for vulnerabilities in 990 cancer cell lines, allows the identification of genes with similar functions or pathways^22, 23^. Among the top correlated genes, FBXL4 showed a strongest positive correlation with PPTC7 (**Fig. 1A**). Notably, FBXL4 and PPTC7 are found to coexist within a co-essential module based on the genetic co-dependency dataset (**Fig. 1B**). Previous study has revealed that the tissues of PPTC7 knockout (KO) mice have markedly diminished mitochondrial content^24^. Considering the striking similarity in mitochondrial defects between PPTC7 knockout (KO) and FBXL4 KO mice^24, 25^, we conducted an investigation to determine whether PPTC7 regulates mitophagy via BNIP3/3L.

**Figure 1.**
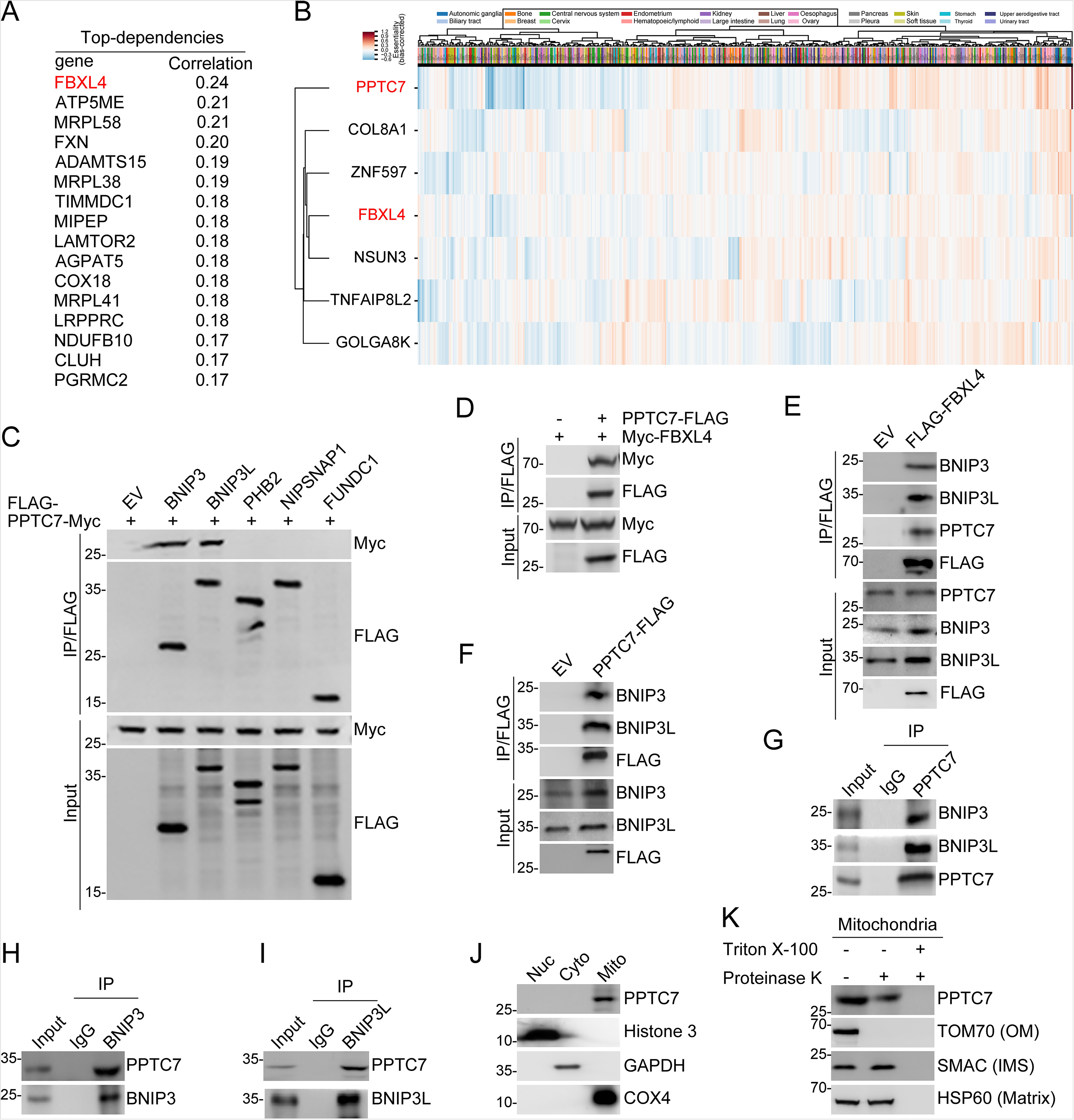
Identification of BNIP3/3L and FBXL4 as PPTC7 interacting proteins. (A, B) DepMap’s data on differences in cell survival resulting from CRISPR genome-wide knockouts. Top-dependencies of PPTC7 are shown in Table (A). (C-F) Western blots (WB) of indicated proteins in input and co-IP samples of anti-FLAG antibody obtained from 293T cells transfected with indicated plasmids. (G-I) WB analysis of the WCL and immunoprecipitates from 293T cells immunoprecipitated with IgG, PPTC7(G), BNIP3(H) and BNIP3L(I) antibody. (J) The cytoplasmic, mitochondrial, and nuclear fractions from HeLa cells were prepared as described in the “Materials and Methods” section. PPTC7 in the three fractions was detected by WB. Histone 3 (nucleus), GAPDH (cytoplasm), and COX4 (mitochondria) were used as subcellular fraction markers. (K) Mitochondria were fractionated from HeLa cells and incubated with (+) or without (-) proteinase K. To disrupt mitochondrial integrity, Triton X-100 was added to the digestion buffer. The indicated proteins in the reactions were detected by WB.

To validate our hypothesis, we first conducted co-immunoprecipitation (co-IP) assays and demonstrated that ectopically-overexpressed PPTC7 interacted with both BNIP3 and BNIP3L. In contrast, PPTC7 did not interact with other mitophagy cargo receptors like PHB2, NIPSNAP1, and FUNDC1, which underscores the highly specific interaction between PPTC7 and BNIP3/3L (**Fig. 1C**). Exogenous co-IP assay results also demonstrated an interaction between PPTC7 and FBXL4 (**Fig. 1D**). Additionally, FLAG-FBXL4 immunoprecipitated endogenous BNIP3/3L and PPTC7 (**Fig. 1E**), and FLAG-PPTC7 immunoprecipitated endogenous BNIP3/3L and FBXL4 (**Fig. 1F**). Finally, we demonstrated that PPTC7 interacted with BNIP3/3L at endogenous levels (**Fig. 1G-I**).

Given that BNIP3/3L and FBXL4 are all outer mitochondrial membrane proteins, we investigated whether PPTC7 was also located on the outer mitochondrial membrane. We further separated the nuclear, mitochondrial, and cytoplasmic fractions of HeLa cells using density-gradient centrifugation methods. PPTC7 is predominantly localized in the mitochondria (**Fig. 1J**), consistent with previous studies reported^20^. Then, we incubated the isolated mitochondrial fractions with proteinase K in the presence or absence of the detergent Triton X-100. Proteins associated with the outer mitochondrial membrane are expected to be protease-sensitive, whereas internal proteins are degraded only after disruption of the mitochondrial membrane by Triton X-100. We found that HSP60 (a mitochondrial matrix protein) and SMAC (a mitochondrial intermembrane space protein) were resistant to proteinase K, whereas TOMM70 (a mitochondrial outer membrane protein) was completely degraded. When proteinase K was added to the mitochondria fraction in the presence of Triton X-100, all the tested proteins were degraded (**Fig. 1K**). As to PPTC7, we noted that proteinase K moderately reduced PPTC7 protein levels, whereas PPTC7 was completely degraded when Triton X-100 was added (**Fig. 1K**). which indicated that a proportion of PPTC7 is located on the outer mitochondrial membrane.

Collectively, these data indicate that PPTC7 specifically interacts with BNIP3/3L on the outer membrane of the mitochondria.

### PPTC7 controls BNIP3/3L protein stability in a protein phosphatase activity-independent manner

Although PPTC7 interacts with FBXL4, PPTC7 protein levels were comparable between parental and FBXL4 KO HeLa cells, indicating that FBXL4 does not affect PPTC7 protein stability (**Supplementary Fig. 1A**). To examine whether PPTC7 regulates BNIP3/3L protein stability in a manner like FBXL4, we utilized siRNA-mediated knockdown (KD) or CRISPR/Cas9-mediated KO to deplete PPTC7 expression in HeLa cells. As shown in **Fig. 2A**, depletion of PPTC7 resulted in a substantial elevation in the steady-state levels of endogenous BNIP3/BNIP3L. Additionally, there was a notable decline observed in the marker proteins within the submitochondrial compartments, such as TOM70 and VDAC1 in the outer membrane, COX4 in the inner membrane, and HSP60 and GRP75 in the matrix. Apart from mitochondria, BNIP3/BNIP3L are also found on peroxisomes where they play a crucial role in promoting pexophagy^26^. Pexophagy is a specific form of autophagy that selectively targets peroxisomes, and it is vital for maintaining the homeostasis of peroxisomes^27^. We observed that the depletion of PPTC7 did not reduce the protein levels of peroxisomal marker proteins (Catalase and PMP70) (**Fig. 2A**), indicating that FBXL4 does not play a role in regulating pexophagy. The impacts of PPTC7 on the protein levels of BNIP3/3L and other mitochondrial marker proteins were also observed in CCF-RC1 and Caki-1 cells (**Supplementary Fig. 1B, C**).

**Figure 2.**
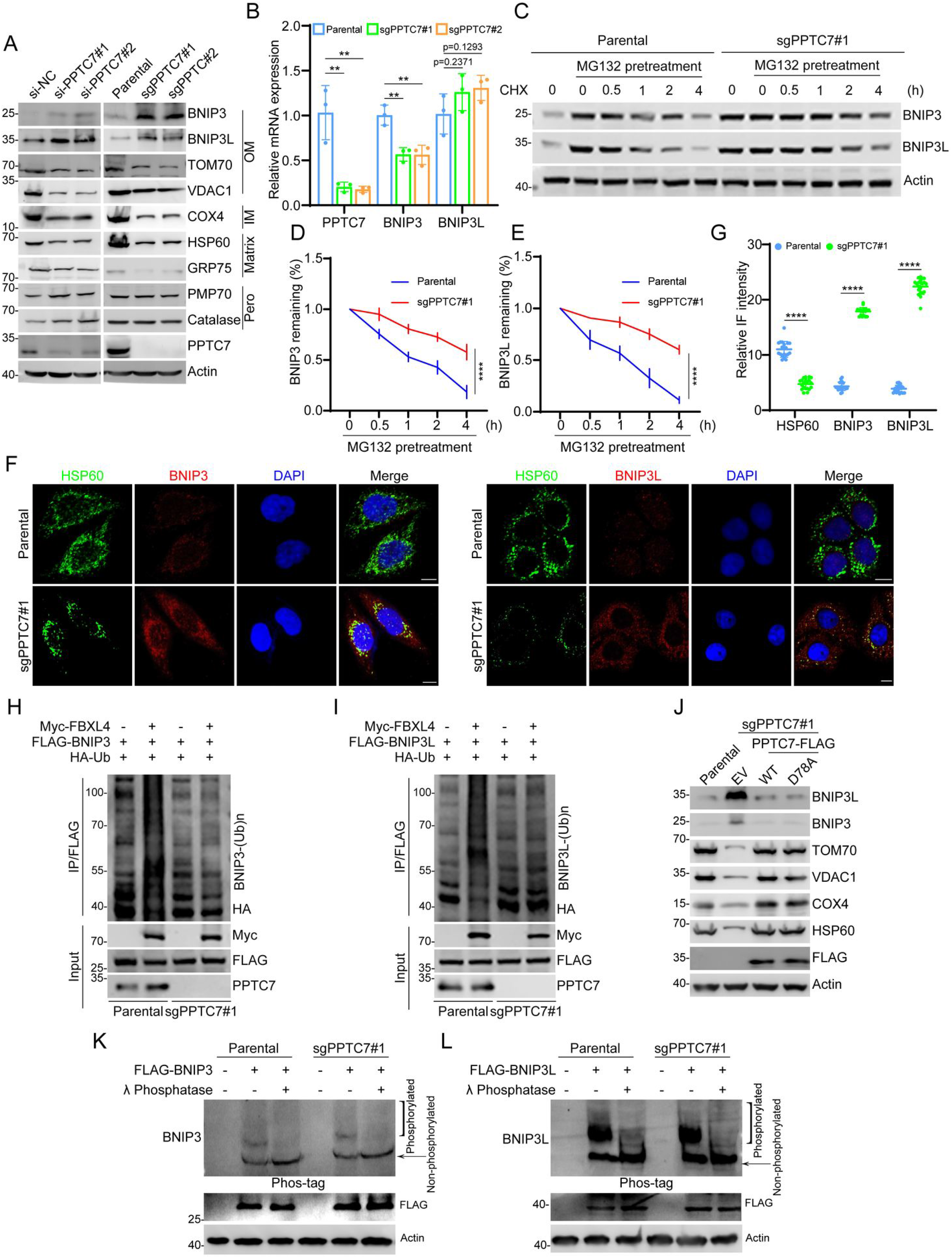
PPTC7 controls BNIP3/3L protein stability in a protein phosphatase activity-independent manner. (A) The WCL from HeLa cells transfected with PPTC7-specific siRNA or negative control (siNC) were prepared for WB with the indicated antibodies. PPTC7 KO cell lines were generated through LentiCRISPRv2 methods. The WCL from parental and PPTC7 KO HeLa cells were prepared for WB with the indicated antibodies. (B) RT qPCR measurement of PPTC7, BNIP3, and BNIP3L mRNA levels in HeLa cells and PPTC7 KO HeLa cells. Data are shown as means ± SD (n = 3). *P* values are calculated by the Two-way ANOVA test. \*\**P* < 0.01. (C-E) WB analysis of the indicated proteins in the WCL of parental and PPTC7 KO HeLa cells pretreated with DMSO or MG132(20 μM) for 5 h and then treated with cycloheximide (CHX, 50 μg/ml) and harvested at different time points. At each time point, the intensity of BNIP3 (D) and BNIP3L (E) was normalized to the intensity of Actin and then to the value at 0 h. *P* values are calculated by the Two-way ANOVA test. \*\*\*\**P* < 0.0001. (F, G) Representative IF images from parental and PPTC7 KO HeLa cells, stained with BNIP3 (or BNIP3L), HSP60 and DAPI. Scale bar, 10 μm. The relative intensity of HSP60, BNIP3 and BNIP3L were quantified and shown in G. Data were shown as means ± SD (n = 20). *P* values are calculated by the Two-way ANOVA test. \*\*\*\**P* < 0.0001. (H, I) WB of the products of in vivo ubiquitination assays from parental and PPTC7 KO HeLa cells transfected with the indicated plasmids. (J) WB analysis of the indicated proteins in WCL from parental and PPTC7 KO HeLa cells stably overexpressing EV, PPTC7-WT-FLAG and PPTC7-D78A-FLAG mutant. (K, L) Parental and PPTC7 KO HeLa cells transfected with the indicated plasmids were lysed, and treated with (+) or without (-) lambada phosphatase (λ phosphatase). These samples were then analyzed by 50□μM Mn^2+^-PhosTag SDS-PAGE (upper panel) or conventional SDS-PAGE (lower panel) and immunoblotted with anti-FLAG antibody.

The mRNA levels of BNIP3 and BNIP3L were either decreased or remained unchanged in PPTC7-depleted cells when compared to the control cells (**Fig. 2B**). This result indicated that the upregulation of BNIP3/3L proteins upon PPTC7 depletion is not achieved through the upregulation of their mRNA levels. Moreover, the half-life of BNIP3/BNIP3L was remarkably prolonged in PPTC7 KO cells (**Fig. 2C-E**). Immunofluorescence (IF) analysis further revealed that the intensity of BNIP3/BNIP3L was markedly upregulated in PPTC7 KO cells, whereas the intensity of mitochondrial marker HSP60 was markedly decreased (**Fig. 2F, G**). Consistently, the endogenous ubiquitination levels of BNIP3/BNIP3L were markedly reduced in PPTC7 KO cells (**Fig. 2H, I**).

PPTC7 has been reported as a protein phosphatase that is localized in the mitochondrial matrix^20, 24^. PPTC7 was reported to dephosphorylate specific mitochondrial proteins, including the mitochondrial import complex protein TIMM50^24^. To assess the significance of PPTC7’s phosphatase activity on BNIP3/BNIP3L and mitochondrial content, we introduced stable overexpression of either PPTC7-WT or its enzymatically inactive D78A mutant into PPTC7 KO cells. Unexpectedly, we observed that the reintroduction of PPTC7-WT or the D78A mutant into PPTC7 KO cells comparably reversed the previously observed accumulation of BNIP3/3L proteins and reduction of mitochondrial proteins caused by PPTC7 deficiency (**Fig. 2J**). To further assess the impact of PPTC7 ablation on the phosphorylation levels of BNIP3/3L, Phos-tag gels were employed to detect changes in the electrophoretic mobility of phosphorylated proteins. We ectopically overexpressed FLAG-BNIP3 in the control or PPTC7 KO cells. We observed BNIP3 bands with higher levels of phosphorylation, and the mobility shifts of the phosphorylated BNIP3 were eliminated after treatment with λ phosphatase. However, the extent of phosphorylated FLAG-BNIP3 was comparable between the control and PPTC7 KO cells (**Fig. 2K**). Similar results were observed when we examined the phosphorylation levels of FLAG-BNIP3L in parental and PPTC7 KO cells (**Fig. 2L).**

Collectively, these data indicate that PPTC7 decreases the protein stability of BNIP3/3L in a manner that is independent of its protein phosphatase activity.

### PPTC7 deficiency activates basal mitophagy in a BNIP3/3L-dependent manner

To examine whether the accumulations of BNIP3/3L proteins are responsible for the reduction of mitochondrial content, we performed individual or combined KD of BNIP3/3L in PPTC7 KO cells. Our results showed that KD of either BNIP3 or BNIP3L partly restored the downregulation of mitochondrial marker proteins caused by PPTC7 KO, whereas the combined KD of BNIP3/3L completely restored it (**Fig. 3A**). In ATG7-KO HeLa cells, we observed a marked increase in BNIP3/BNIP3L protein levels upon PPTC7 KD, while the levels of mitochondrial marker proteins showed no obvious change. These results indicated that PPTC7 plays a crucial role in regulating mitochondrial content through the canonical autophagy pathway (**Supplementary Fig. 1D**). To monitor mitophagy, we utilized Mthagy Dye, which accumulates in intact mitochondria and exhibits weak fluorescence under normal conditions. Upon induction of mitophagy, damaged mitochondria fuse with lysosomes, leading to strong fluorescence emission from Mthagy Dye^28^. We found that PPTC7 KO significantly enhanced mitophagy signals. KD of BNIP3 or BNIP3L alone partially blocked the activation of mitophagy, whereas combined KD of BNIP3/3L completely blocked it (**Fig. 3B, C**). Additionally, we evaluated several mitochondrial metabolic indexes to assess mitochondrial functions. PPTC7 KO resulted in decreased rates of oxygen consumption (OCR) and ATP production, along with an increase in lactate production. Notably, combined KD of BNIP3/BNIP3L effectively reversed these effects (**Fig. 3D-F**).

**Figure 3.**
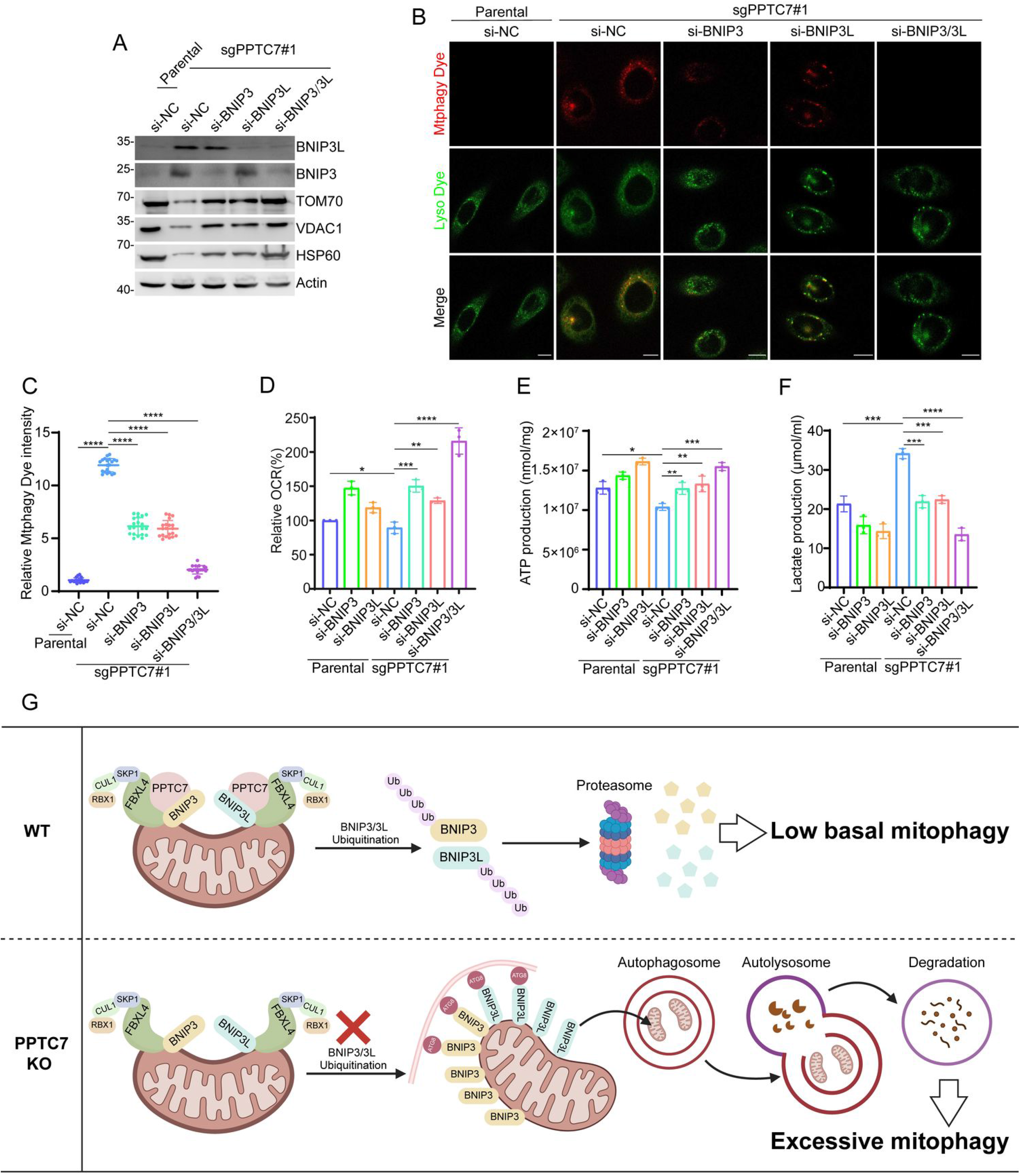
PPTC7 deficiency activates mitophagy, which is dependent on BNIP3/BNIP3L. (A) WB analysis of the indicated proteins in the WCL from parental or PPTC7 KO HeLa cells transfected with indicated siRNAs. (B, C) Representative IF images from parental and PPTC7 KO HeLa cells transfected with indicated siRNAs, stained with Mtphagy Dye and Lyso Dye. Scale bar, 10 μm. The relative intensity of Mtphagy Dye was quantified and shown in (C). Data were shown as means ± SD. (n = 20). *P* values are calculated by the One-way ANOVA test. \*\*\*\**P* < 0.0001. (D) The OCR of parental and PPTC7 KO HeLa cells transfected with the indicated siRNA or siNC were measured using OCR assay Kit. Data are shown as means ± SD (n = 3). (E) The intracellular ATP production of parental and PPTC7 KO HeLa cells transfected with indicated siRNAs were measured using ATP production assay Kit. Data are shown as means ± SD (n = 3). (F) The intracellular lactate levels of parental and PPTC7 KO HeLa cells transfected with indicated siRNAs were measured using Lactate Assay Kit. Data are shown as means ± SD (n = 3). *P* values are calculated by the One-way ANOVA test and the Two-way ANOVA test in (D–F). \**P* < 0.05; \*\**P* < 0.01; \*\*\**P* < 0.001; \*\*\*\**P* < 0.0001.

Collectively, these data indicate that PPTC7 deficiency induces hyperactive mitophagy due to BNIP3/3L accumulation.

### PPTC7 deficiency activates BNIP3/3L-dependent mitophagy in the *Pptc7* KO mice models

To gain a better understanding of the pathophysiological roles of PPTC7 on BNIP3/3L proteins and mitophagy *in vivo*, we generated *Pptc7* knockout mice models (**Fig. 4A**). Consistent with a previous study showing that *Pptc7* KO in mice caused fully penetrant lethality, no mice survived at P10 in our experimental settings^24^. Thus, we collected multiple tissues, including brain, heart, liver, muscle, and intestine from day 20 mice. WB analysis indicated that the tissues of *Pptc7^-/-^* mice exhibited varying degrees of upregulation in BNIP3/3L protein levels, compared to those of wild-type (WT) mice. Additionally, mitochondrial proteins were inevitably downregulated across all the examined tissues (**Fig. 4B**).

**Figure 4.**
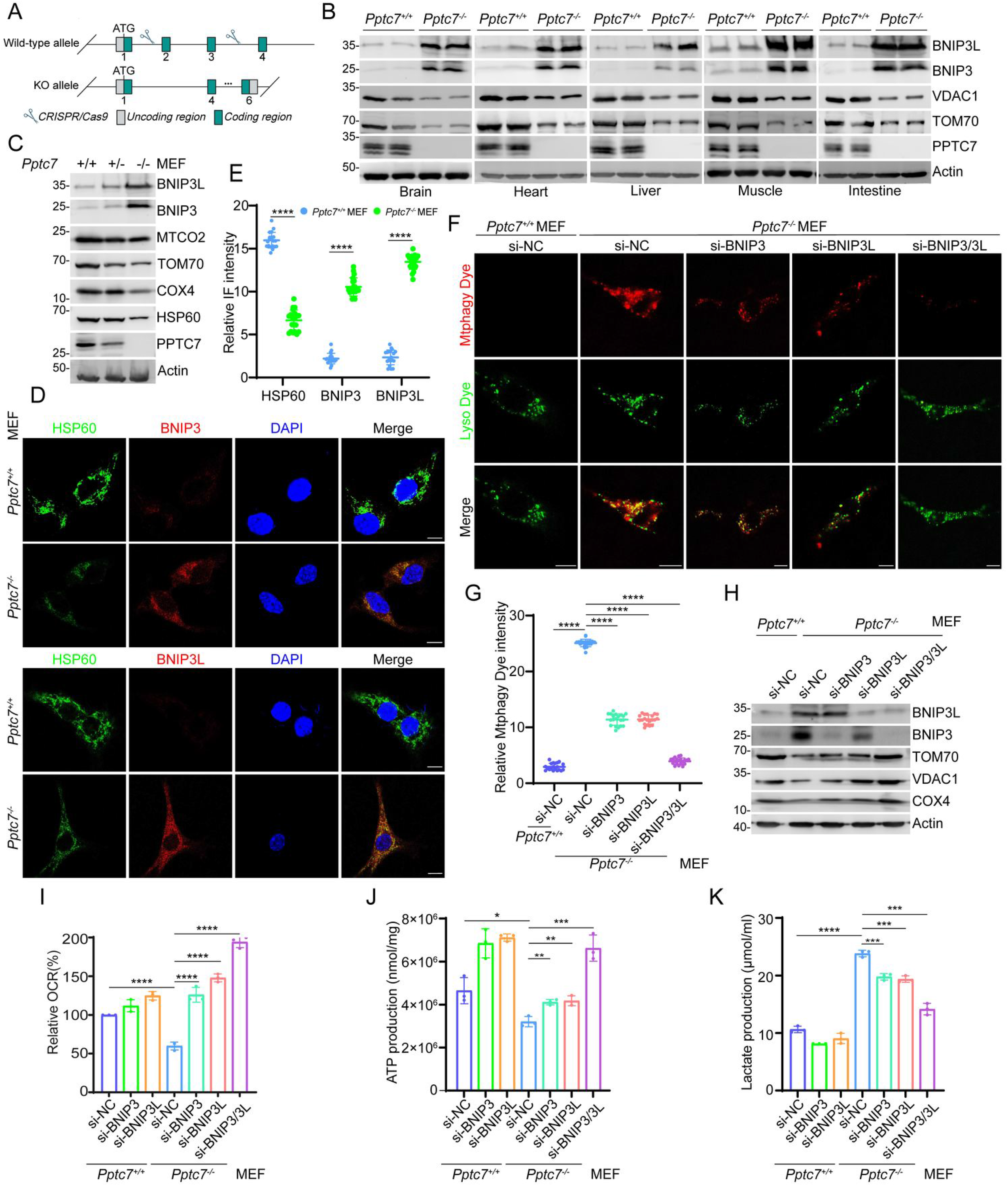
BNIP3/BNIP3L-dependent mitophagy was abnormally activated in *Pptc7* KO mice models. (A) Strategy to generate *Pptc7* KO mice using CRISPR/Cas9 methods. (B) WB analysis of the indicated proteins in WCL from the indicated tissues of *Pptc7^+/+^* and *Pptc7^-/-^* mice. (C) WB analysis of the indicated proteins in WCL from *Pptc7^+/+^*, *Pptc7^+/-^* and *Pptc7^-/-^* MEFs. (D, E) Representative IF images from *Pptc7^+/+^* and *Pptc7^-/-^* MEFs, stained with BNIP3 (or BNIP3L), HSP60 and DAPI. Scale bar, 10 μm. The relative intensity of HSP60, BNIP3 and BNIP3L were quantified and shown in (E). Data were shown as means ± SD (n = 20). *P* values are calculated by the Two-way ANOVA test. \*\*\*\**P* < 0.0001. (F, G) Representative IF images from *Pptc7^+/+^* and *Pptc7^-/-^* MEFs transfected with the indicated siRNAs, stained with Mtphagy Dye and Lyso Dye. Scale bar, 10 μm. The relative intensity of Mtphagy Dye was quantified and shown in (D). Data were shown as means ± SD (n = 20). *P* values are calculated by the One-way ANOVA test. \*\*\*\**P* < 0.0001. (H) WB analysis of the indicated proteins in WCL from *Pptc7^+/+^* and *Pptc7^-/-^* MEFs transfected with the indicated siRNAs. (I) The OCR of *Pptc7^+/+^*and *Pptc7^-/-^* MEFs transfected with indicated siRNAs were measured using OCR assay Kit. Data are shown as means ± SD (n=3). (J) The intracellular ATP production of *Pptc7^+/+^* and *Pptc7^-/-^*MEFs transfected with the indicated siRNA or siNC were measured using ATP production assay Kit. Data are shown as means ± SD (n=3). (K) The intracellular lactate levels of *Pptc7^+/+^* and *Pptc7^-/-^* MEFs transfected with the indicated siRNAs were measured using Lactate Assay Kit. Data are shown as means ± SD (n=3). *P* values are calculated by the One-way ANOVA test and the Two-way ANOVA test in (I-K). \**P* < 0.05; \*\**P* < 0.01; \*\*\**P* < 0.001; \*\*\*\**P* < 0.0001.

We next prepared mouse embryonic fibroblasts (MEFs). In homozygous *Pptc7^-/-^* MEFs, BNIP3/3L protein levels were markedly elevated, while mitochondrial marker protein levels were reduced compared to WT or heterozygous Pptc7^+/-^ MEFs (**Fig. 4C-E**). *Pptc7^-/^*^-^ MEFs exhibited a high basal level of basal mitophagy, which was almost undetectable in WT MEFs. Notably, KD of BNIP3 or BNIP3L alone partially inhibited Pptc7 KO-induced mitophagy activation, while combined KD of BNIP3/3L completely blocked it (**Fig. 4F-H**). Furthermore, *Pptc7^-/-^* MEFs displayed decreased rates of OCR and ATP production, along with an increase in lactate production. Importantly, these effects were largely reversed by combined KD of BNIP3/3L (**Fig. 4I-K**).

Collectively, these data indicate that PPTC7 deficiency in mice induces hyperactive mitophagy due to BNIP3/3L accumulation.

### PPTC7 is an essential co-factor of the SCF^FBXL4^ E3 ubiquitin ligase complex

As PPTC7 does not contain any domain ubiquitin-proteasome system, it does not seem likely to directly promote the ubiquitination and degradation of BNIP3/3L. The aforementioned results indicated that depletion of PPTC7 resulted in BNIP3/3L accumulation, resembling the phenotypes observed in FBXL4-KO cells. Since PPTC7 interacts with FBXL4, we investigated whether PPTC7 acts as a co-factor of the SCF^FBXL4^ E3 ubiquitin ligase complex to regulate BNIP3/3L protein stability. Surprisingly, similar to the outcomes from the PPTC7 depletion experiments, we observed that the exogenous overexpression of PPTC7-WT or the catalytically inactive D78A mutant in parental HeLa cells both markedly increased the steady-state levels of endogenous BNIP3/3L and simultaneously reduced mitochondrial content (**Fig. 5A, Supplementary Fig. 2A, B**). The mRNA levels of BNIP3/3L were either reduced or remained unchanged in PPTC7-overexpressed cells (**Fig. 5B**). These results underscored the critical role of maintaining precise control over PPTC7 protein levels in regulating the abundance of BNIP3/3L. In contrast, ectopic overexpression of PPTC7-WT in FBXL4-KO cells showed no impact on the protein levels of BNIP3/3L and mitochondrial content (**Fig. 5C**). IF analysis also revealed that the overexpression of PPTC7-WT or the D78A mutant in FBXL4-KO cells failed to reverse the BNIP3/3L accumulation caused by FBXL4 deficiency (**Fig. 5D, E**).

**Figure 5.**
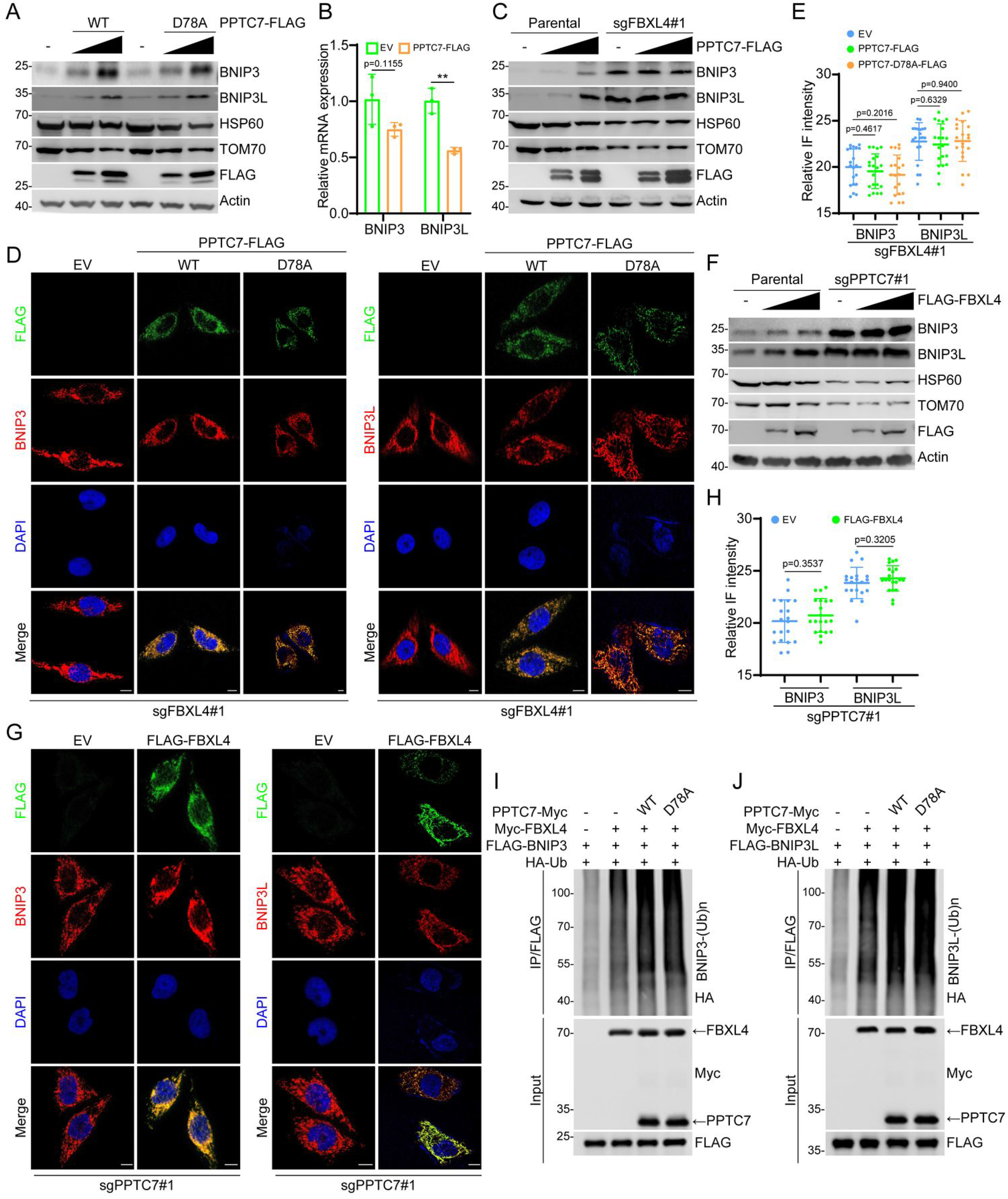
PPTC7 is an essential co-factor of SCF^FBXL4^ E3 ubiquitin ligase complex. (A) WB analysis of the indicated proteins in WCL from parental HeLa cells stably overexpressing EV, PPTC7-WT-FLAG, or PPTC7-D78A-FLAG mutant. (B) RT-qPCR measurement of BNIP3 and BNIP3L mRNA levels in HeLa cells stably overexpressing EV or PPTC7-FLAG. Data are shown as means ± SD (n = 3). *P* values are calculated by the Two-way ANOVA test. \*\**P* < 0.01. (C) WB analysis of the indicated proteins in WCL from FBXL4 KO HeLa cells stably overexpressing EV or PPTC7-FLAG. (D-E) Representative IF images from FBXL4 KO HeLa cells stably overexpressing EV, PPTC7-WT-FLAG, or PPTC7-D78A-FLAG mutant, stained with BNIP3 (or BNIP3L), HSP60, and DAPI. Scale bar, 10 μm. The relative intensity of HSP60, BNIP3, and BNIP3L were quantified and shown in (E). Data were shown as means ± SD (n = 20). *P* values are calculated by the Two-way ANOVA test. (F) WB analysis of the indicated proteins in WCL from PPTC7 KO HeLa cells stably overexpressing EV or FLAG-FBXL4. (G, H) Representative IF images from PPTC7 KO HeLa cells stably overexpressing EV or FLAG-FBXL4, stained with BNIP3 (or BNIP3L), HSP60, and DAPI. Scale bar, 10 μm. The relative intensity of HSP60, BNIP3, and BNIP3L were quantified and shown in (H). Data were shown as means ± SD (n = 20). *P* values are calculated by the Two-way ANOVA test. (I, J) WB of the products of in vivo ubiquitination assays from 293T cells transfected with the indicated plasmids.

Similarly, ectopic overexpression of FBXL4 in parental HeLa cells markedly increased the steady-state levels of endogenous BNIP3/3L and simultaneously reduced mitochondrial content (**Fig. 5F, Supplementary Fig. 2C**). However, ectopic overexpression of FBXL4 in PPTC7 KO cells did not affect the protein levels of BNIP3/3L and mitochondrial content (**Fig. 5G, H**). Finally, we found that co-expression of either PPTC7-WT or the D78A mutant could further enhance the FBXL4-induced ubiquitination of BNIP3 or BNIP3L (**Fig. 5I, J**).

Collectively, these data suggest that PPTC7 act as an essential co-factor of the SCF^FBXL4^ E3 ubiquitin ligase complex to facilitate the ubiquitination and degradation of BNIP3/3L.

## DISCUSSION

Using i*n vitro* cell culture and *Pptc7* knockout mice models, we demonstrate that PPTC7 acts as an essential co-factor of the SCF^BFXL4^ complex to facilitate ubiquitin-proteasomal degradation of BNIP3/3L, thereby keeping basal mitophagy at a very low level (**Fig. 3G**). This active regulation may allow a rapid mitophagy response under certain conditions, through disrupting SCF^BFXL4/PPTC7^-mediated BNIP3/3L degradation. However, the upstream molecular events and physiological functions underlying these are still elusive.

Our results also corroborated by a recent study showing that *Pptc7* KO mice caused hyperactive mitophagy in a BNIP3/3L-dependent manner^29^. During the preparation of this manuscript, a published study also reported that PPTC7 acts as a mitophagy sensor to control BNIP3/3L degradation to regulate mitophagy^29^. This study furthermore demonstrated that Pptc7 KO-induces perinatal lethality can be rescued by Bnip3L KO, indicating Bnip3L-mediated mitophagy may be a critical downstream event in *in vivo*^30^. One unexpected phenomenon we observed is that overexpression of PPTC7 also resulted in the stabilization of BNIP3/3L and induction of mitophagy, similar to the phenotypes seen in PPTC7 knockout. It might be possible that the subunit stoichiometry is a critical determinant of the correct function of the SCF^FBXL4/PPTC7^ complex, although the molecular details warranted further investigation. It also suggests that simply upregulating the protein levels of PPTC7 is insufficient to suppress BNIP3/3L-dependent mitophagy; Both PPTC7 and FBXL4 might need to be co-regulated.

PPTC7 is reported as a mitochondrial matrix phosphatase in yeast and mammalian cells. using submitochondrial fraction method, we demonstrated that a proportion of PPTC7 is located on the outer mitochondrial membrane, it explains why PPTC7 can interact with outer mitochondrial membrane protein FBXL4 and BNIP3/3L. We also showed that reintroduction of PPTC7-WT or its enzymatically inactive mutant into PPTC7 KO cells comparably reversed the accumulation of BNIP3/3L proteins. Moreover, the phosphorylation levels of BNIP3/3L were comparable between parental and PPTC7 KO cells as judged by phos-tag gels. These results indicated that the phosphatase activity may be dispensable for its role in regulating BNIP3/3L turnover. PPTC7 has a homologous protein in yeast, but FBXL4 does not have a homologous protein in yeast. An unpublished study in a master’s thesis showed that RNAi knockdown of CG12091 (the homolog PPTC7 protein of Drosophila) resulted in drastic changes in mitochondrial morphology and neurodegeneration in Drosophila. Furthermore, overexpression of CG12091-WT or the enzymatically inactive mutant CG12091mu significantly impeded the development of the Drosophila compound eye and abnormal mitochondria were observed in CG12091-WT or CG12091mu overexpressed photoreceptor cells^31^. From an evolutionary perspective, we speculate that SCF^BFXL4/PPTC7^-mediated mitophagy regulation may emerge after yeast but before Drosophila.

Similar to PPTC7, several metabolic enzymes have the ability to function as modulators of E3 ubiquitin ligases or deubiquitinases. For instance, guanosine 5′-monophosphate synthetase (GMPS) interacts with deubiquitinase USP7 to promote Histone H2B deubiquitination by USP7. However, the activation of USP7 by GMPS is not directly associated with the GMP synthesis reaction, indicating that its role in modulating ubiquitination operates through a mechanism distinct from its enzymatic activity^32^. Serine hydroxymethyltransferase 2 (SHMT2), a metabolic enzyme responsible for converting serine to glycine and a tetrahydrofolate-bound one-carbon unit, serves as a subunit of the BRISC deubiquitinase complex, which plays various roles in DNA damage repair and immune regulation. Notably, the SHMT2 bound to BRISC is catalytically inactiv^33^. CRL3^KLHL10^ E3 ubiquitin ligase complex depends on succinyl-CoA synthetase β subunit (A-Sβ), a Krebs cycle enzyme, to ubiquitylate mitochondrial membrane proteins^34^. Thus, these proteins, including PPTC7, can be classified as moonlighting proteins which perform multiple autonomous and often unrelated functions without partitioning these functions into different domains of the protein^35^.

## MATERIALS AND METHODS

### Cell culture

293T, HeLa, and MEF cells were cultured in Dulbecco’s Modified Eagle’s Medium (DMEM) supplemented with 10% fetal bovine serum (FBS). Caki-1 and CCF-RC1 cells were cultured in Roswell Park Memorial Institute 1640 Medium (RPMI-1640) supplemented with 10% FBS. All cells were cultured at 37□°C under 5% CO_2_ in a humidified incubator. DNA fingerprinting and PCR were performed to verify the authenticity of the cell lines and to ensure they are free of mycoplasma infection. Transient transfection was conducted using EZ Trans (Shanghai Life-iLab Biotech) or Lipofectamine 8000 (Beyotime). For lentiviral transfection, pLVX overexpression plasmids and virus-packing constructs were transfected into 293T cells. The viral supernatant was collected after 48 h. The cells were then infected with the viral supernatant in the presence of polybrene (8 µg/ml) and selected in growth media containing puromycin (1.5 μg/ml). The sequences of gene-specific siRNAs are listed in **Supplementary Table 1**.

### Antibodies and chemicals

The information of antibodies and chemicals used in this study is listed in **Supplementary Table 2, 3**.

### Gene KO cell line generation

To knockout *PPTC7*, *FBXL4*, or *ATG7* genes in human cells, CRISPR/Cas9 protocols were employed. The sgRNAs were designed using an online CRISPR design tool (http://crispr.mit.edu) and subcloned into the LentiCRISPRv2 vector from Dr. Feng Zhang’s lab. Guide RNA containing the target sequence and virus-packing constructs were transfected into 293T cells. The viral supernatant was collected after 48 h. HeLa, Caki-1, or CCF-RC1 cells were then infected with the viral supernatant in the presence of polybrene (8 µg/ml). The cells were selected with puromycin (2 µg/ml) for 3-7 days. Successful KO of each gene was verified via Western blot (WB) analysis. The sequences of gene-specific sgRNAs are listed in **Supplementary Table 1**.

### *In vivo* ubiquitination assays

For ubiquitination analysis, HA-tagged ubiquitin and other indicated plasmids were co-transfected into 293T or HeLa cells. 36□h After transfection, MG132 was added to the medium for 4–6□h before harvesting. Cells were collected, lysed, and boiled in 1% SDS buffer (Tris-HCl, pH 7.5, 0.5□mM EDTA, 1□mM DTT) for 10□min. Immunoprecipitation was performed in 10-fold diluted lysates with 0.5% NP-40 buffer, and the ubiquitination levels of BNIP3 or BNIP3L were detected via WB.

### RNA isolation and quantitative real-time reverse transcription PCR (qRT-qPCR)

Total RNAs were isolated from cells using the TransZol Up reagent (TRANS) following the manufacturer’s instructions. Concentrations and purity of RNAs were determined by measuring the absorption of ultra-violet lights using a NanoDrop spectrophotometer (Thermo). cDNAs were reversed-transcribed using a HiScript III RT SuperMix for qPCR (Vazyme), followed by amplification of cDNA using ChamQ SYBR qPCR Master Mix (Vazyme). The relative mRNA levels of genes were quantified using the 2-ΔΔCt method, with normalization to Actin. The sequences of primers are listed in **Supplementary Table 1**.

### IF and confocal microscopy

HeLa or MEF were seeded on glass coverslips in 12-well plates and harvested at 80% confluence. The cells were washed with PBS and fixed with 4% paraformaldehyde in PBS at RT for 30□min. After permeabilization with 0.1% Triton X-100 for 30 min and then in the blocking solution (PBS plus 5% donkey serum), for 1h at room temperature (RT). The cells were then incubated with primary antibodies at 4°C overnight. After washing with PBST buffer, fluorescence-labelled secondary antibodies were applied. DAPI was utilized to stain nuclei. The glass coverslips were mounted on slides and imaged using a confocal microscope (LSM880, Zeiss) with a 63*/1.4NA Oil PSF Objective. Quantitative analyses were performed using ImageJ software.

### Isolation of nucleus, cytoplasm, and mitochondria

HeLa cells were prepared for nuclear, cytoplasmic, and mitochondrial extraction by density-gradient centrifugation. Briefly, 5□×□10^6^ HeLa cells were washed three times with PBS and then suspended by using hypotonic solution (140□mM KCl, 10□mM EDTA, 5□mM MgCl_2_, 20□mM HEPES (pH 7.4), and the protease inhibitor). Then, the cells were ground with a glass homogenizer in an ice bath for 25 strokes. Nuclear, cytoplasmic, and mitochondrial fractions were separated through differential centrifugation (800 ×g, 10□min, 4□°C and 12,000×g, 35□min, 4□°C). The supernatant (cytoplasmic fraction) and pellet (mitochondrial fraction) were collected, and the pellet was further washed with wash buffer (800□mM KCl, 10□mM EDTA, 5□mM MgCl_2_, and 20□mM HEPES (pH 7.4), and the protease inhibitor) for three times and yield the final mitochondrial fraction. To confirm that pure extracts were obtained, the mitochondrial, nuclear, and cytoplasmic fractions were separated by SDS-PAGE, and the presence of mitochondrial COX4, nuclear Histone H3, and cytoplasmic GAPDH was detected via WB.

### Mitochondrial protein localization assays

Mitochondria were purified using the methods described above. The control group was left untreated. The second group was treated with proteinase K (3□μM). The third group was treated with proteinase K (3□μM) and 0.5% Triton-X-100 solution. Three groups of samples were placed in a 37□°C water bath for 30□min. The samples were prepared and then detected by WB.

### Oxygen consumption assays

OCR was measured under basal conditions in the presence of the mitochondrial inhibitor oligomycin (0.25 μM, Calbiochem) at 37°C. OCR was calculated by the oligomycin-induced changes in comparison to basal rates. HeLa and MEF cells were seeded at a density of 2×10^4^ cells in the cell culture microplate. The total protein of each well was determined by Bradford assay and used as the reference to normalize the OCR.

### ATP measurement assays

HeLa or MEF cells were seeded in 6-well plates at 2×10^6^ cells per well and cultured overnight. Then, the cells were transfected with the indicated siRNAs. 48 h after transfection, the cells were collected for determination of ATP production using the ATP assay kit (Beyotime).

### Lactate measurement assays

HeLa or MEF Cells were seeded into 6-well plates at a density of 1×10^5^ cells per well and cultured overnight. According to the manual of the lactate acid assay kit (Solarbio), the resulting color was measured at 570 nm using a microplate absorbance reader (Bio-Rad).

### Generation and breeding of *Pptc7* Cas9-KO mice

*Pptc7* CRISPR/Cas9-KO mice were designed and generated by GemPharmatech Co., Ltd. In brief, Cas9 mRNA was *in vitro* transcribed with mMESSAGE T7 Ultra Kit (Ambion) according to the manufacturer’s instructions, and subsequently purified using the MEGAclearTM kit (Thermo) Cas9 sgRNA was *in vitro* transcribed using the MEGAshortscript kit (Thermo) and subsequently purified using MEGAclearTM kit. The transcribed Cas9 mRNA and sgRNA as well as a 200 base pairs single-stranded oligo deoxy nucleotide (ssODN) were co-injected into zygotes of C57BL/6J mouse. Obtained F0 mice were validated by PCR and Sanger sequencing. The F0 mice with expected point mutation were chosen and crossed with C57BL/6J mice to produce F1 mice. Genotyping was performed by PCR analysis of tail DNA. The sequences of sgRNAs are listed in **Supplementary Table 1**. Mice were maintained under a 12 h/12 h light/dark cycle at 22-25°C and 40–50% humidity with standard food and water available ad libitum. All animal procedures were conducted in accordance with the animal care committee of Fudan University.

### MEFs generation and immortalization

Timed pregnant female mice at embryonic day 12.5 to 14.5 were sacrificed, and the embryos were carefully dissected to remove the cerebrum, internal organs, and limbs. The remaining tissues were cut into small pieces and treated with trypsin-EDTA (0.25%) for 10 min at 37°C. The trypsin was neutralized with DMEM, a complete medium supplemented with 10% FBS and 1% penicillin/streptomycin. The culture media were changed every 2-3 days until the cells reached confluence. To immortalize MEFs, they were passaged up to approximately 10 times before infection with lentiviral vectors expressing the SV40 large T-antigen. Stable transduction was achieved with puromycin selection. The successful integration of the immortalizing gene was confirmed through Sanger sequencing and WB analysis.

### Statistical analysis

Band intensities of WB results were calculated by ImageJ in accordance with the manufacturer’s instructions. Statistical analysis was performed using Prism 8.0 (GraphPad Software, Inc., San Diego, CA, USA) and Excel (Microsoft Corp., Redmond, CA, USA). Pooled results were expressed as the mean□±□SEM. Comparisons between groups were made via One-way analysis of variance (ANOVA) or Two-way ANOVA. Statistical significance was set at *P*□≤□0.05; ns no significance; \**P*□<□0.05; \*\**P*□<□0.01; \*\*\**P*□<□0.001; \*\*\*\**P*□<□0.0001.

## Acknowledgments

The image in graphical model is produced by Biorender (https://app.biorender.com/gallery).

## Author contributions

C.W. and S.M. conceived the study. X.X. and Y.C. performed the experiments and data analyses. C.W., S.M., and Y.L. analyzed and interpreted the data. C.W. wrote the manuscript.

## Data and materials availability

All data needed to evaluate the conclusions in the paper are present in the paper and/or the Supplementary Materials.

