## Supplementary Materials for "PPTC7 acts as an essential co-factor of the SCF^FBXL4^ ubiquitin ligase complex to restrict BNIP3/BNIP3L-dependent mitophagy"

Figures. S1 to S2

**Other Supplementary Materials for this manuscript include the following:**

Table S1 to S3

**Supplementary Table 1.** Sequence information.

**Supplementary Table 2.** Antibody information.

**Supplementary Table 3.** Cell cultures, chemicals, and kits.

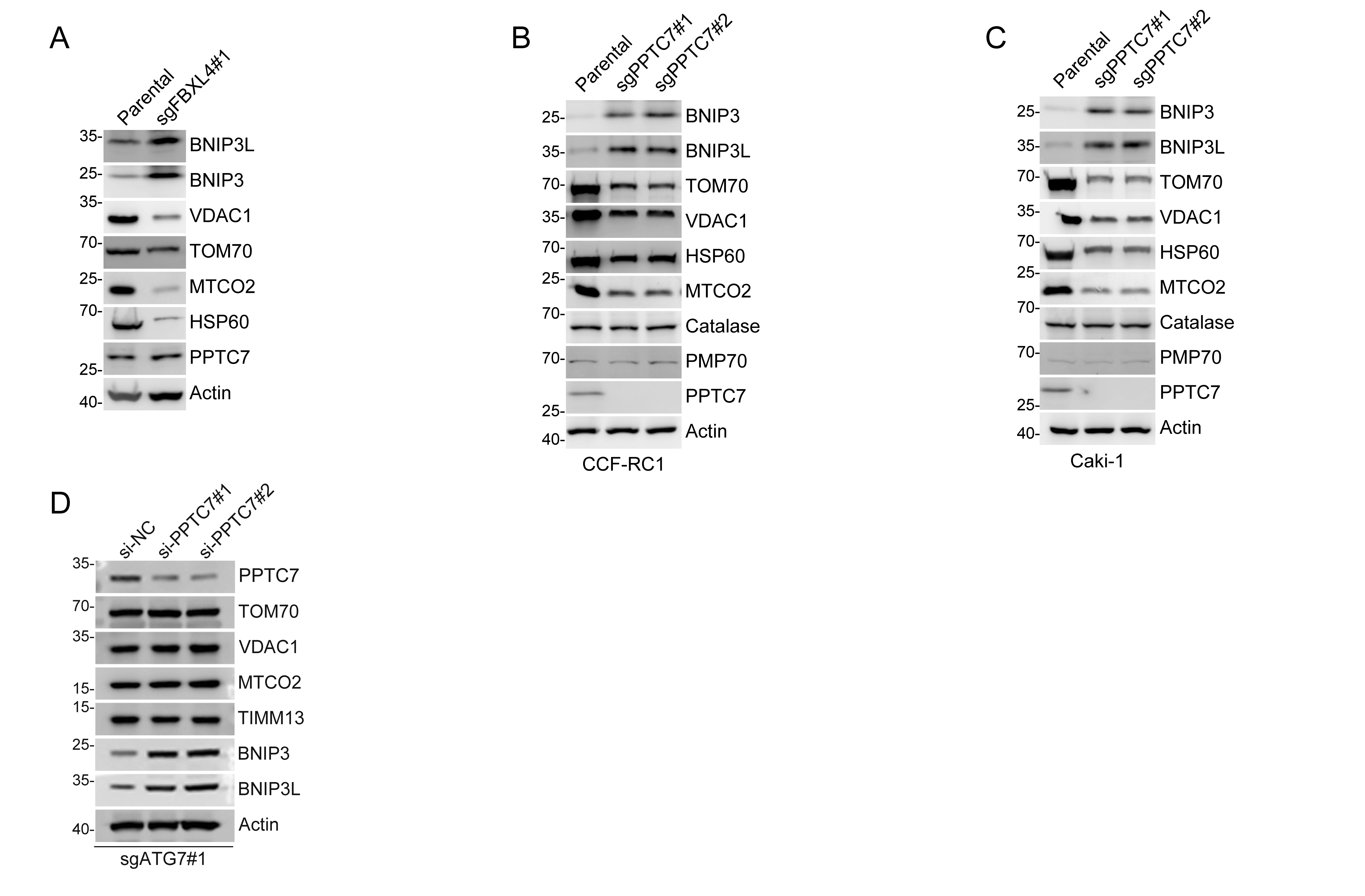

**Supplementary Figure 1. PPTC7 controls BNIP3/3L protein stability in a protein phosphatase activity-independent manner (related to Figure. 2, 3).**

(A) WB analyses of the indicated proteins in the WCL from parental and FBXL4-KO HeLa cells.

(B, C) WB analyses of the indicated proteins in the WCL from parental and PPTC7-KO CCF-RC1 (B) or Caki-1 (C) cells.

(D) WB analyses of the indicated proteins in the WCL from ATG7-KO HeLa cells transfected with PPTC7-specific siRNAs or siNC.

**Supplementary Figure 2. PPTC7 is an essential co-factor of SCF^FBXL4^ E3 ubiquitin ligase complex (related to Figure 5).**

(A, B) Representative IF images from HeLa cells overexpressing the indicated plasmids, stained with BNIP3 (or BNIP3L), HSP60, and DAPI. Scale bar, 10 μm. The relative intensity of HSP60, BNIP3, and BNIP3L was quantified and shown in (B). Data were shown as means ± SD (n=20).

(C, D) Representative IF images from parental HeLa cells overexpressing the indicated plasmids, stained with BNIP3 (or BNIP3L), HSP60, and DAPI. Scale bar, 10 μm. The relative intensity of HSP60, BNIP3, and BNIP3L was quantified and shown in (D). Data were shown as means ± SD (n=20).

*P* values are calculated by the Two-way ANOVA test in (B, D). *****P* <0.0001.
